# BenchMHC: a reproducible framework for peptide-MHC presentation prediction

**DOI:** 10.64898/2026.10.22.760998

**Authors:** Théomé Borck, Marion Chabrol, Daniel Colombo, Arthur Imbert, Martin Perier-Dulhoste, Konstantin Volzhenin, Maren Lang, Nicolas Lopez Carranza

## Abstract

Predicting which peptides an MHC molecule presents is the first step in neoantigen and epitope discovery pipelines. Most published predictors are released for inference only, and each is trained on a different dataset, so the effect of a new method cannot be separated from the effect of its training data. We present **BenchMHC**, an open framework that reproduces, retrains and evaluates pMHC presentation models on public data only, and that records every implementation decision made in a reproduction. Our NetMHCpan-4.1 reproduction reaches 0.794 per-allele Mean Top-K against 0.790 for the official webserver, and 0.0025 Median FRANK against 0.0022. It scores 280,000 peptide/allele pairs in 27 s, about 350 times faster than the original binary. We then examine the training data. In the public NetMHCpan-4.1 eluted-ligand sets, 86 to 99% of hits are found in the human proteome, against about 12% of their decoys, so a model can separate hits from decoys using this difference rather than features related to MHC binding. Redrawing the decoys within the source protein of each hit and matching their cysteine status, with no other change, raises per-allele Mean Top-K on the regenerated `v2.0.0` evaluation set from 0.734 to 0.755, and increases Median FRANK on CD8 epitopes from 0.0025 to 0.0053. The same change improves one evaluation set and degrades the other. Code (Apache-2.0) and model checkpoints (CC BY-NC-4.0): https://github.com/instadeepai/BenchMHC-public.

## 1 Introduction

MHC molecules bind short peptides and display them on the cell surface, where T cells can inspect them. Predicting which peptides are presented for a given allele is the first step in pipelines that select peptides for immunological use: tumour neoantigen discovery for personalised cancer immunotherapy [11, 10], T-cell epitope identification for pathogens, vaccine design, autoimmunity studies and transplantation matching. Predictors are trained on two kinds of data: in-vitro binding-affinity (BA) assays, and mass-spectrometry eluted-ligand (EL) data, which is either single-allelic (SA) or multiallelic (MA). NetMHCpan-4.1 [9] is the most widely cited pan-allele MHC-I predictor. Recent work adds expression features [3], transformer architectures [12] or improved pan-allele generalisation [7].

Most of these models are released for inference only, and each is trained on a different dataset. The contribution of an architecture therefore cannot be separated from the contribution of its training data. When a model reports better numbers than a baseline on its own benchmark, there is no shared setup in which to test whether the method or the data is responsible. Users who want to retrain a model have to re-curate decoys, re-calibrate percentile ranks and re-implement MA deconvolution themselves.

### Contributions

(i) **BenchMHC**, an open Python framework for reproducing, retraining and evaluating pMHC presentation models on public data only, in which each reproduced decision is recorded as identical to the original, under-specified in the source publication, or deliberately changed (§2). (ii) A reproduction of NetMHCpan-4.1 that matches the official webserver on the two metrics commonly reported for this task, and is about two orders of magnitude faster at inference (§3). (iii) An analysis of the public training data, which contains a species imbalance between hits and decoys and a cysteine-depletion artefact. Correcting both, without changing the architecture, improves one evaluation set and degrades the other (§4). We report both results.

## 2 The BenchMHC framework

BenchMHC is written in Python 3.12. It uses Polars [13] for lazy, streaming operations over the millions of (peptide, allele, decoy) rows that NetMHCpan-scale training requires, PyTorch [8] for the network, and PyTorch Lightning [5] for the training loop. Separate subpackages hold the Click command-line interface, a Polars-backed `DataModule`, the feature-engineering layer, custom layers, losses and callbacks, and shared utilities for MA deconvolution, percentile-rank calibration and BLAST. Figure 1 shows the pipeline; Appendix A lists the full set of commands.

**Figure 1:**
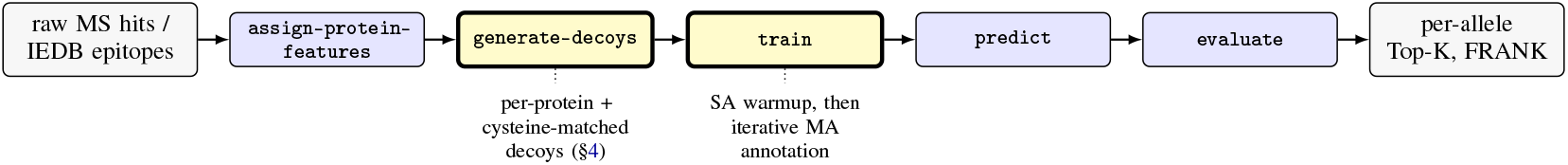
The BenchMHC pipeline: a chain of `bench-mhc` subcommands from raw massspectrometry hits to per-allele metrics. Grey boxes are data files and blue boxes are commands; the two yellow boxes are the stages this paper examines, decoy construction (§4) and the iterative multi-allelic annotation loop (§3). Optional stages (peptide refinement, core precomputation, percentile-rank calibration, plotting) are omitted; see Appendix A.

### Recording implementation decisions

For each reproduced method we record every implementation decision and label it 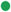 identical to the original, 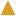 not fully specified in the source publication and reconstructed by us, or 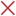 changed on purpose. NetMHCpan-4.1 gives 20 such decisions (Table 1): nine identical to the original, seven not fully specified, two changed on purpose, and two that depend on which dataset version is used. For the under-specified rows the published descriptions do not fix a value, and we used the most direct reading of the text in each case. The two deliberate changes are that we do not balance EL against BA examples within a batch, and that we validate on SA data only, where the original mixes SA and MA. A reader can check each row, change the ones they disagree with, and rerun the pipeline.

**Table 1:** Implementation decisions in the BenchMHC NetMHCpan-4.1 reproduction, each labelled 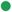 identical to the original, 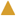 not fully specified in the published literature and reconstructed by us, or 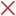 changed on purpose. Every choice in the reproduction is documented and justified in the model documentation page released with the code.

| Implementation decision | Status |
| --- | --- |
| NNAlign technique (9-mer cores via indels) | ● |
| Ensemble of 50 MLPs (10 seeds × 5 folds) | ● |
| Hidden size of the MLP (55 vs 56/66) | ▲ |
| MLP weight initialisation | ▲ |
| Input features (BLOSUM62, pseudo-sequence, flanks, lengths) | ● |
| BLOSUM62 amino-acid encoding (exact matrix variant) | ▲ |
| Dual heads (binding affinity, eluted ligand) | ● |
| Instance-based max-pool over 9-mer cores | ● |
| 9-mer core selection for the BA head | ▲ |
| Loss function (BCE for EL + MSE for BA) | ▲ |
| Optimiser (SGD, lr = $5 \times 10^{-3}$) | ● |
| Number of epochs (200) | ● |
| Early-stopping patience | ▲ |
| Batch size | ▲ |
| Batch balancing (EL:BA ≈ 1:1) | ✕ |
| Iterative annotation on MA data (NNAlign_MA) | ● |
| Prediction-score rescaling (z-score with schedule) | ● |
| Mixing SA and MA in the validation set | ✕ |
| Training data (original release) | ● / ✕ <sup>†</sup> |
| Evaluation data (original release) | ● / ✕ <sup>†</sup> |

<sup>†</sup> ● for the v0.0.0 reproduction (Sec. 3); ✕ for the v1.1.0 re-curation (Sec. 4).

## 3 Reproducing NetMHCpan-4.1

NetMHCpan-4.1 is an ensemble of 50 MLPs (10 seeds *×* 5 folds) trained with NNAlign [4, 2]. Each MLP takes a flat encoding of a 9-mer core, the 34-residue HLA pseudo-sequence, insertion and deletion lengths, flank counts and peptide-length indicators, all encoded with BLOSUM62 [6]. Peptides of length *k* ≠ 9 are mapped to candidate cores by inserting X tokens (*k <* 9) or deleting a window of residues (*k >* 9); a max-pool over cores passes the gradient through the highest-scoring core only. Two heads predict IC_50_ (MSE loss) and presentation likelihood (BCE loss), with unlabelled entries masked. Training uses SGD with a constant learning rate of 5 *×* 10^−3^, batch size 1024, and at most 200 epochs with early stopping. MA configurations follow NNAlign_MA: 20 warmup epochs on SA data, then repeated rescale, deconvolute and fine-tune steps. Appendix B gives the architecture diagrams and the rescaling schedule.

### Comparison with the official tool

Table 2 compares our reproduction, trained on the original NetMHCpan-4.1 data, with the official webserver on two metrics: per-allele Mean Top-K on MS `ligands v0.0.0` (45,416 MHC-I eluted ligands, 36 HLA alleles) and per-epitope Median FRANK on CD8 epitopes (1,660 IEDB epitopes validated in T-cell assays). Top-K is the positive predictive value at *K*, with *K* set to the number of true ligands of that allele, so it is the fraction of the allele’s *K* top-scoring peptides that are ligands; FRANK is the fraction of the peptides of an epitope’s source protein that score above the epitope, so 0 is a perfect ranking [9]. The EL head reaches 0.794 Mean Top-K against 0.790, and 0.0025 Median FRANK against 0.0022. Figure 2 shows that the per-allele distributions overlap. Because all data configurations are in one codebase and use the same splits, we can also compare models trained on different subsets of the data. EL data gives the largest contribution on MS ligands, while BA data alone already performs well on CD8 epitopes (Appendix C).

**Table 2:** Main results. Per-allele Mean Top-K on the MS ligands evaluation sets (higher is better) and per-epitope Median FRANK on the CD8 epitopes set (lower is better). The BenchMHC blocks use the same architecture, optimiser, schedule, ensemble size and random-seed grid, and differ only in training data. “n/a”: the official webserver was not re-scored on the regenerated `v2.0.0` set. Boldface marks the best EL-score number in each column. Mean FRANK and per-configuration results are in Appendix C.

| Model (training data) | Head | Mean Top-K ↑ | | Median FRANK ↓ |
| --- | --- | --- | --- | --- |
|  |  | MS v0.0.0 | MS v2.0.0 | CD8 epitopes |
| Official NetMHCpan-4.1 | BA | 0.754 | n/a | 0.0030 |
| Official NetMHCpan-4.1 | EL | 0.790 | n/a | **0.0022** |
| BenchMHC v0.0.0 (original) | BA | 0.755 | 0.701 | 0.0030 |
| BenchMHC v0.0.0 (original) | EL | **0.794** | 0.734 | 0.0025 |
| BenchMHC v1.1.0 (re-curated) | BA | 0.744 | 0.692 | 0.0029 |
| BenchMHC v1.1.0 (re-curated) | EL | 0.733 | **0.755** | 0.0053 |

**Figure 2:**
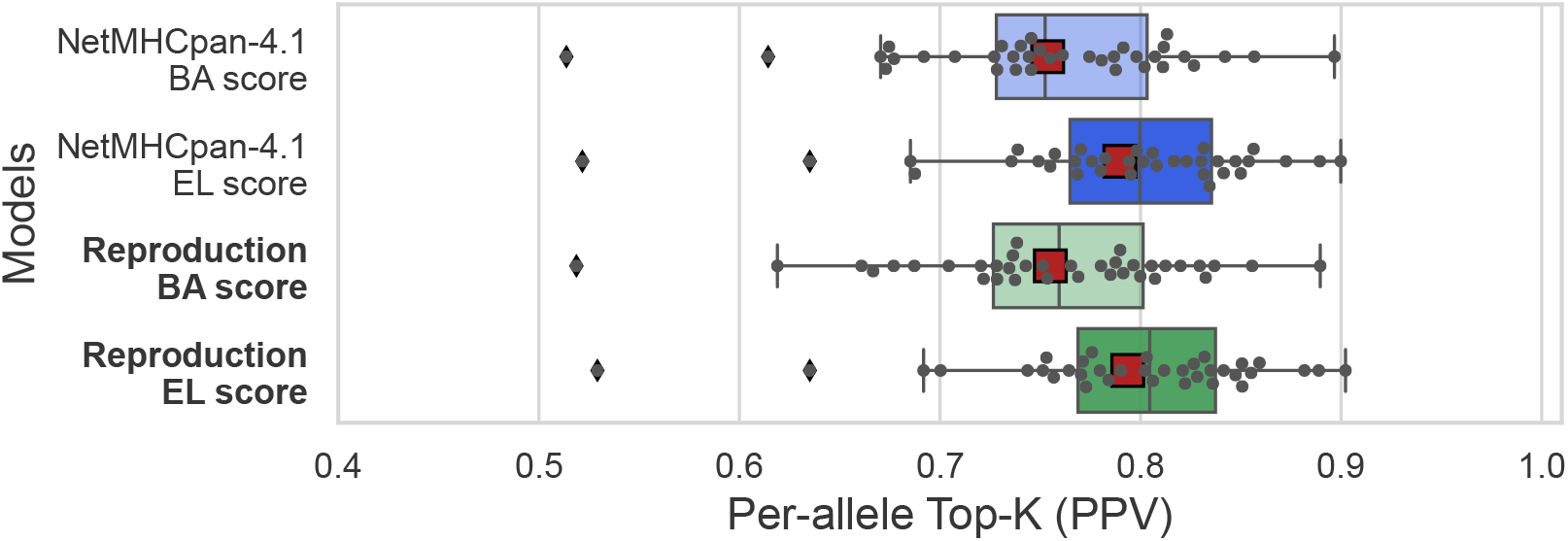
Per-allele Top-K (PPV) on the original `MS ligands` MHC-I evaluation set, for the official NetMHCpan-4.1 and for our reproduction, both heads. Each box is the distribution over the 36 alleles; dots are individual alleles and the red square is the mean. Data-type ablations are in Appendix C.

### Inference speed

Two changes reduce inference time without altering the model or its outputs. First, predict prepares the scoring data once and reuses a single trainer for all 50 members, so feature preparation runs once instead of 50 times. Second, a fused ensemble groups members by architecture signature. The BLOSUM62 features and 9-mer cores do not depend on the model, so they are computed once per batch, and each group’s stacked layers are applied as batched `einsum` products. This replaces 50 sequential forward passes with a few batched ones. On one NVIDIA P100 the 280,000-pair reference set is scored in about 27 s, about 350 times faster than the original single-process binary and about 40 times faster than a 10-process parallel wrapper. Scores agree with the per-member path to about 10^−6^ (Appendix D).

## 4 Decoy construction in the public training data

The reproduction gives access to the full data path, so we can examine the training data directly. Two properties of the public NetMHCpan-4.1 data are described below. Neither is a modelling choice.

i. **Species imbalance between hits and decoys**. The positive EL hits come mostly from human immunopeptidomics. The decoys are natural peptides drawn from UniProt-SwissProt, which is mostly non-human, and they are not matched to the source organism of the hit. Figure 3 shows the size of the difference: 86 to 99% of SA and MA hits are found in the human proteome, against about 12% of their decoys. A model can therefore separate hits from decoys using residue statistics that differ between human and non-human protein fragments, instead of features related to MHC binding. IEDB [14] contributes a large number of viral and bacterial epitopes, so this affects a substantial part of the data. The BA peptides are largely synthetic and show no such difference.
ii. **Cysteine depletion**. Mass-spectrometry sample preparation modifies cysteines, usually by alkylation, so Cys-containing peptides are under-represented among EL hits. If decoys are drawn without controlling for cysteine content, the model lowers its scores for Cys-containing peptides in general. This reflects the assay rather than binding.

**Figure 3:**
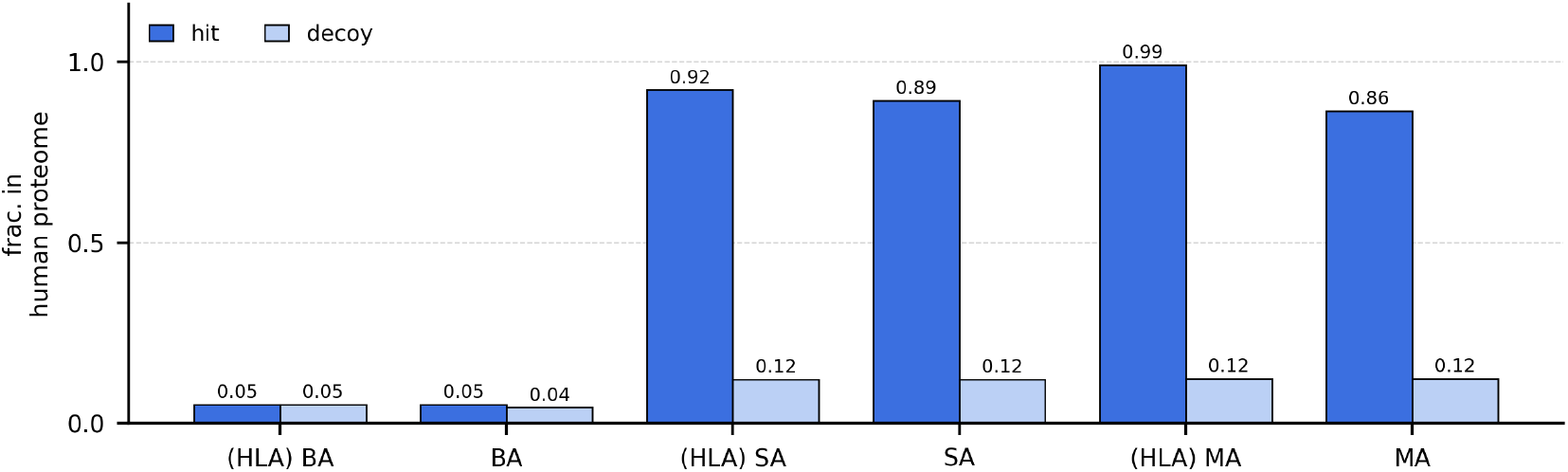
Fraction of unique training peptides found in the proteome associated with the hg38 human genome, split into hits and decoys, per data-type configuration of the original NetMHCpan-4.1 data. On the eluted-ligand sets (SA, MA) hits are 86 to 99% human and their decoys about 12%. Binding-affinity (BA) peptides are largely synthetic and show no such difference.

### Correction

We change the `generate-decoy`s stage. For each hit (*p*_*h*_, HLA_*h*_) with source protein *π*_*h*_ we (1) restrict the decoy pool to *k*-mers of *π*_*h*_, so that decoys come from the same source protein as the hit, and (2) keep only candidates whose cysteine status matches that of *p*_*h*_, falling back to the opposite status when none is available. All other settings are unchanged: splits, hit set, model, optimiser, schedule, ensemble size and random seeds are identical to the reproduction. Datasets and the models trained on them carry a semantic version (Appendix E); the corrected training data is `v1.1.0` and its matched evaluation set is `v2.0.0`.

### Results

Retraining the same ensemble on `v1.1.0` raises the EL-head per-allele Mean Top-K from 0.734 to 0.755 on the `v2.0.0` evaluation set, an increase of about 3%, with no change to the architecture. The increase is spread across the 36 alleles rather than coming from a few (Figure 4). On the CD8 epitopes set the same models perform worse: Median FRANK increases from 0.0025 to 0.0053 (Table 2). One possible explanation is that the CD8 epitopes are mostly viral, while matching EL decoys to their source proteins, which are mostly human, makes the training data more human-specific.

**Figure 4:**
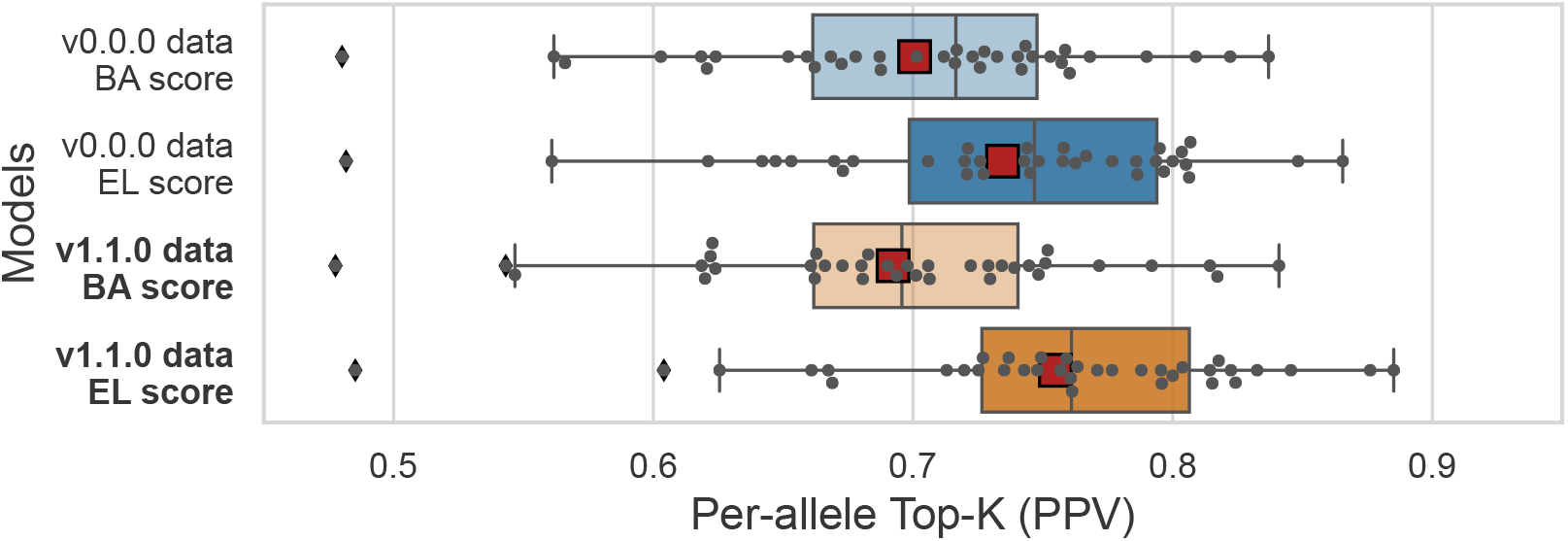
Per-allele Top-K (PPV) on the `v2.0.0` MS-ligands evaluation set, for the ensemble trained on the original `v0.0.0` data (blue) and the one trained on the corrected `v1.1.0` data (orange), both heads. Each box is the distribution over the 36 alleles; dots are individual alleles and the red square is the mean. The `v1.1.0` EL-score distribution is shifted towards higher values across the alleles. Per-allele bars are in Appendix C.

## 5 Discussion

A single change to the decoy sampling, with no change to the model, improves per-allele Mean Top-K on MS ligands and degrades Median FRANK on CD8 epitopes. Reporting only the MS-ligands number would present this as a data-curation improvement. Both numbers come from the same models, so the direction of the result depends on which evaluation set is used. This matters for how pMHC benchmarks are read: the decoy-generating process is part of what a model is scored on, so a change that removes an artefact from one evaluation set can reduce accuracy on another. Measuring this requires regenerating the training data and retraining, which is not possible with an inference-only release.

### Limitations

The species and cysteine corrections were applied together, so we cannot separate their individual effects. The reported numbers are single runs of a 50-MLP ensemble, without seed-level confidence intervals. The `v2.0.0` evaluation set was itself regenerated, so we compare the same model on the same set. We have reproduced one method; comparing architectures on identical data requires at least a second one, and NetMHCIIpan-4.3 and NetMHCpanExp-1.0 [3] are in progress.

## Availability

Data curation, decoy generation, the training loop, calibration and evaluation are shared across models, so adding a method requires a model definition and a configuration file rather than a new pipeline. We welcome external reproductions, datasets and corrections. Appendix F lists what the release contains. Code and documentation: https://github.com/instadeepai/Bench_MHC-public.

## A Pipeline and command surface

Figure 5 shows the full `bench-mhc --help` surface. Seven subcommands form the raw-hits-topredictions chain, of which Figure 1 draws the five that every run uses: a`ssign-protein-features` BLASTs [1] the hits against SwissProt and the human proteome to recover the source protein and flanking residues; `generate-decoys` samples decoys per source protein with cysteine-status matching (§4); `train` fits the model with an SA warmup followed by iterative MA annotation; `calibrate` (optional) computes per-allele percentile ranks against a reference set of 10,000 random 9-mers; and `predict, evaluate` and `generate-performance-plot` produce scores, metrics and plots. The remaining subcommands (peptide refinement, core precomputation, allele-feature formatting) are optional. Each subcommand has a dedicated guide in the repository documentation.

**Figure 5:**
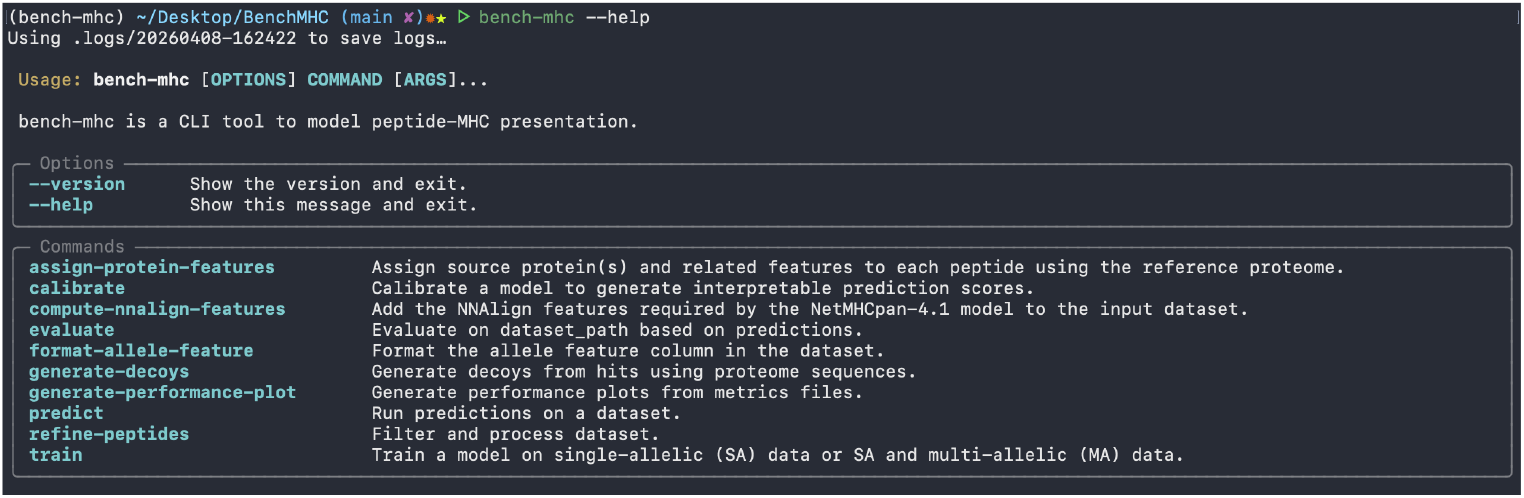
Output of `bench-mhc --help`. The CLI surface exposes the full pipeline; each subcommand has a dedicated guide in the repository documentation.

## B Architecture and training schedule

Figure 6 illustrates NNAlign core generation and Figure 7 the network. The flat 868-dimensional input is fed to a single hidden layer (size 56 or 66 in our reproduction) and a dual head sharing all parameters except the final linear layer.

**Figure 6:**
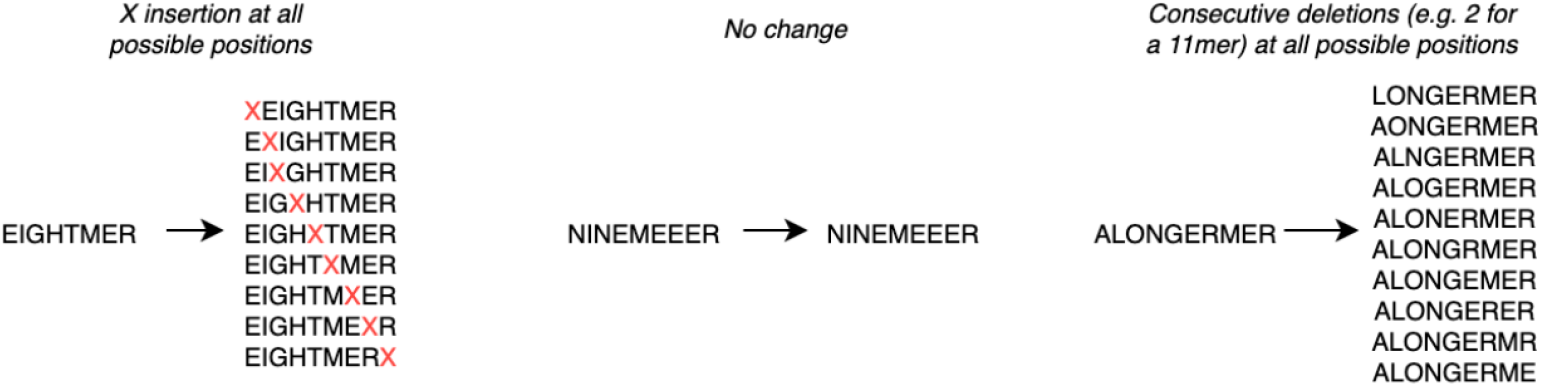
NNAlign preprocessing: every peptide of length *k* ≠ 9 is mapped to a set of candidate 9-mer cores by inserting X tokens (if *k <* 9) or by deleting a window of consecutive residues (if *k >* 9); each core inherits the corresponding insertion/deletion length. Training back-propagates the gradient through the maximum-scoring core only.

**Figure 7:**
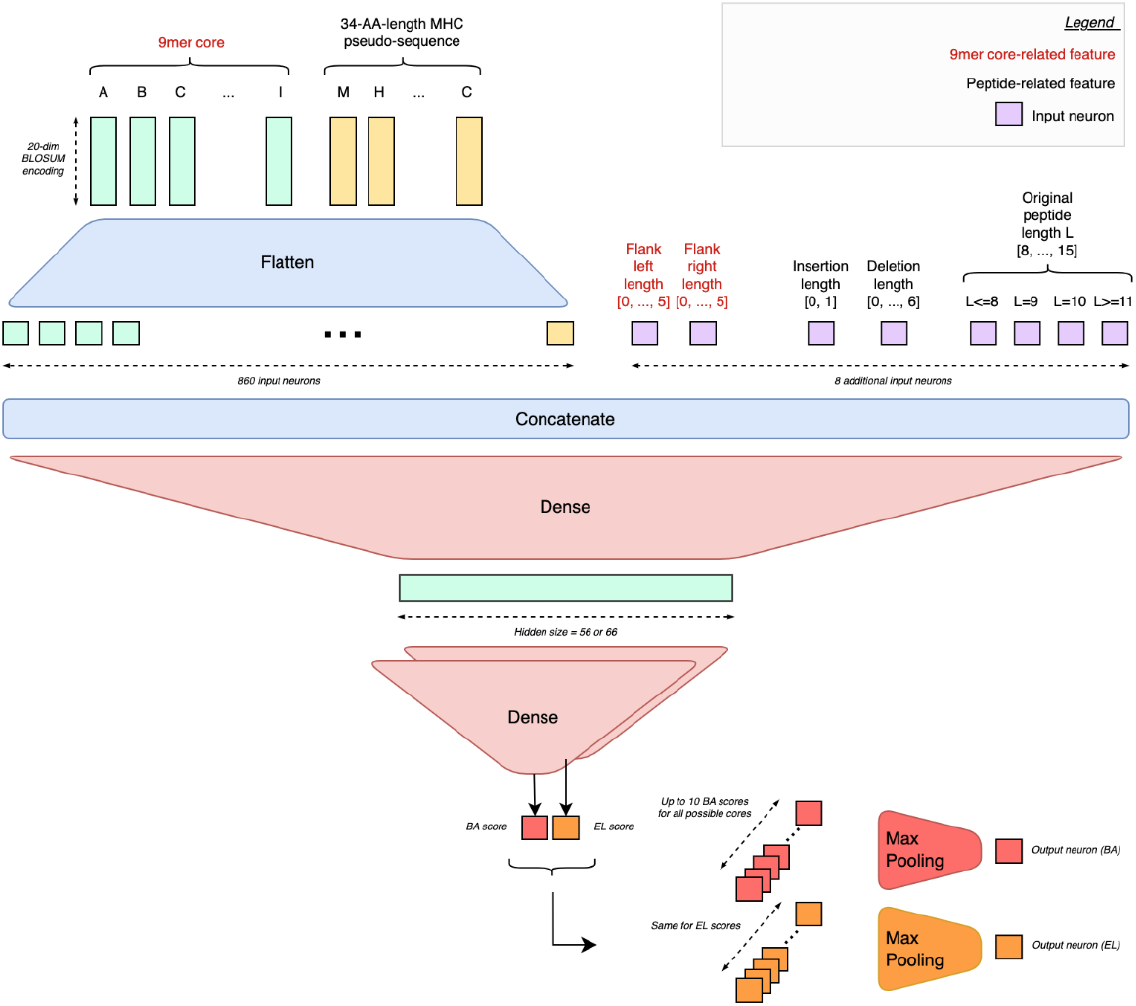
NNAlign architecture. The flat 868-dimensional input is fed to a single hidden layer (size 56 or 66 in our reproduction) and a dual head: the BA head predicts an IC_50_ score via MSE, the EL head predicts the likelihood of ligand presentation via BCE. The two heads share all parameters except the final linear layer.

Training uses SGD with a constant learning rate of 5 *×* 10^−3^, batch size 1024, and at most 200 epochs with early stopping on validation loss (patience 10 to 20 depending on head configuration). SA-only configurations are a standard fit. Configurations including MA data follow the NNAlign_MA [2] recipe: warm up for 20 epochs on SA data, then enter the iterative annotation loop. At each iteration we rescale per-allele scores so they are comparable across alleles, then deconvolute the MA samples by keeping, per multi-allelic bag, the allele with the highest rescaled score. The rescaling normalises each allele’s score statistics (mean and standard deviation, estimated on a reference set with z-score outlier filtering) by interpolating them toward the global pooled statistics with a logistic weight

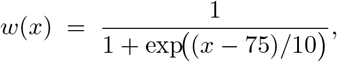

where *x* is the epoch index, so the rescaling moves smoothly from per-allele statistics (*w* ≈ 1, early) to global statistics (*w* ≈ 0) around epoch 75. We then merge with the SA pool and fine-tune for one epoch.

## C Extended metrics and data-configuration ablations

Table 3 repeats Table 2 with the Mean FRANK column. Figures 8 and 9 show per-allele Top-K on the MS ligands set and per-epitope FRANK on the CD8 epitopes set for the official webserver, our full **BA** + **SA** + **MA** reproduction, and every intermediate data-type configuration. A “(HLA)” prefix marks a model trained on HLA (human) data only; an unprefixed label denotes training on all data, HLA and non-HLA. In addition to the comparison in Figure 2, eluted-ligand data accounts for most of the performance on MS ligands, and binding-affinity and non-HLA data each add a smaller amount. On the CD8 epitopes set, binding-affinity data alone already performs well, and adding eluted-ligand and non-HLA data generally improves the scores further, although the effect of non-HLA data is small and not uniform across configurations. Figure 10 gives the per-allele bars underlying Figure 4, for the same two ensembles.

**Table 3:** Full metrics table. Per-allele Mean Top-K on the MS ligands evaluation set (higher is better) and per-epitope Median FRANK Score on the CD8 epitopes evaluation set (lower is better). The BenchMHC blocks share the same architecture, optimiser, schedule, ensemble size and random-seed grid: they differ only in the training data. Boldface marks the best EL-score number per column. “n/a” marks the official webserver, which we did not re-score on the regenerated `v2.0.0` evaluation set.

| Model | Head | Per-allele Mean Top-K ↑ | | Median FRANK ↓ | Mean FRANK ↓ |
| --- | --- | --- | --- | --- | --- |
|  |  | MS lig. v0.0.0 | MS lig. v2.0.0 | (CD8 epit.) | (CD8 epit.) |
| Official NetMHCpan-4.1 * | BA score | 0.754 | n/a | 0.0030 | 0.0114 |
| Official NetMHCpan-4.1 * | EL score | 0.790 | n/a | **0.0022** | **0.0105** |
| BenchMHC v0.0.0 (§3) | BA score | 0.755 | 0.701 | 0.0030 | 0.0117 |
| BenchMHC v0.0.0 (§3) | EL score | **0.794** | 0.734 | 0.0025 | 0.0138 |
| BenchMHC v1.1.0 (§4) | BA score | 0.744 | 0.692 | 0.0029 | 0.0119 |
| BenchMHC v1.1.0 (§4) | EL score | 0.733 | **0.755** | 0.0053 | 0.0335 |

\* Official NetMHCpan-4.1 numbers obtained by running the BenchMHC evaluation pipeline on the public webserver scores; see [9].

**Figure 8:**
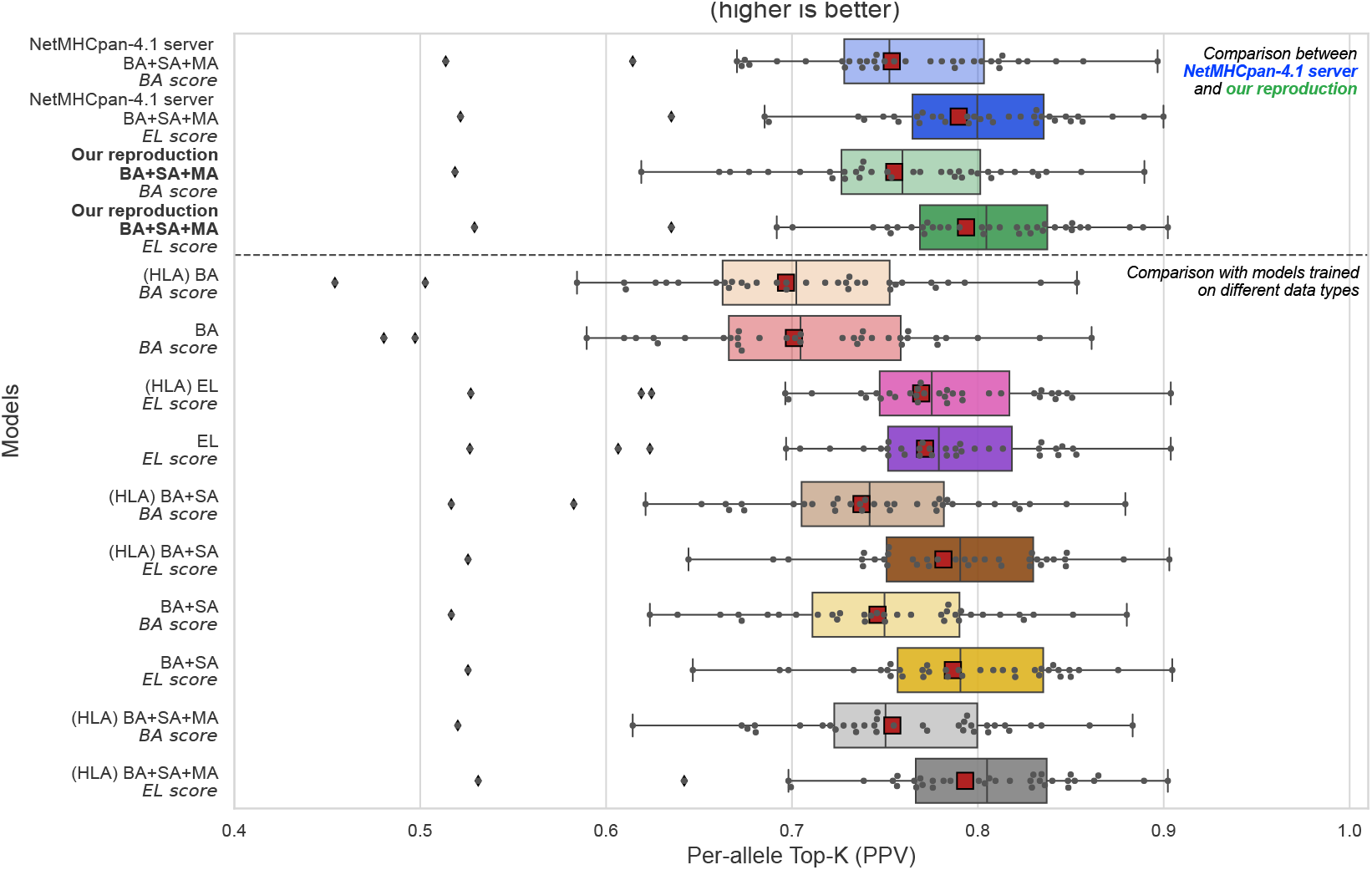
Per-allele Top-K (PPV) on the original MS ligands MHC-I evaluation set (higher is better). Top block: the official NetMHCpan-4.1 webserver versus our **BA** + **SA** + **MA** reproduction (BA-score and EL-score heads). Bottom block: models trained on each data-type configuration. Each box is the distribution over the 36 alleles.

**Figure 9:**
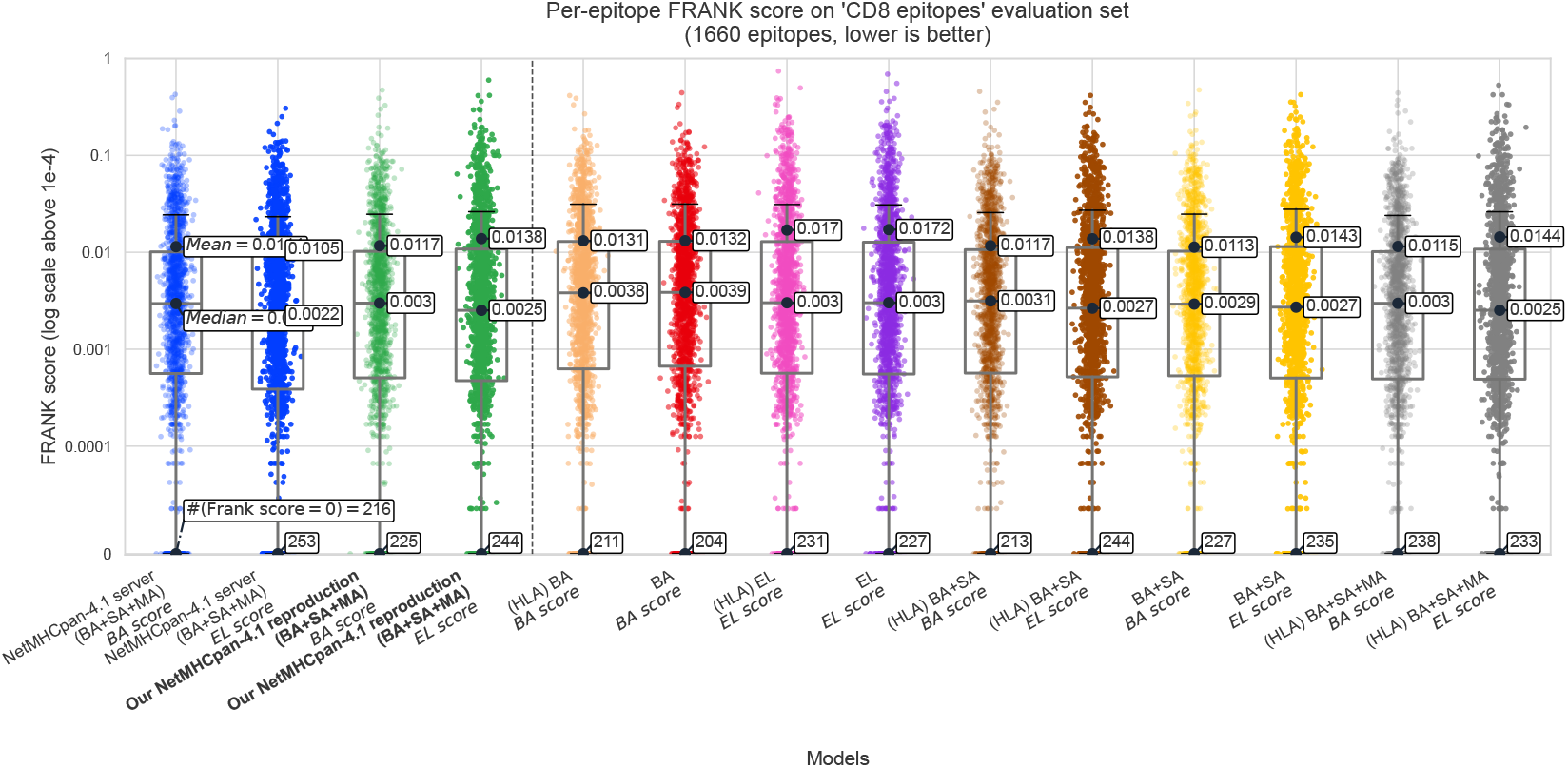
Per-epitope FRANK score on the CD8 epitopes evaluation set (log scale, lower is better) for the same models as Fig. 8. Each point is one epitope; boxes mark the median and quartiles.

**Figure 10:**
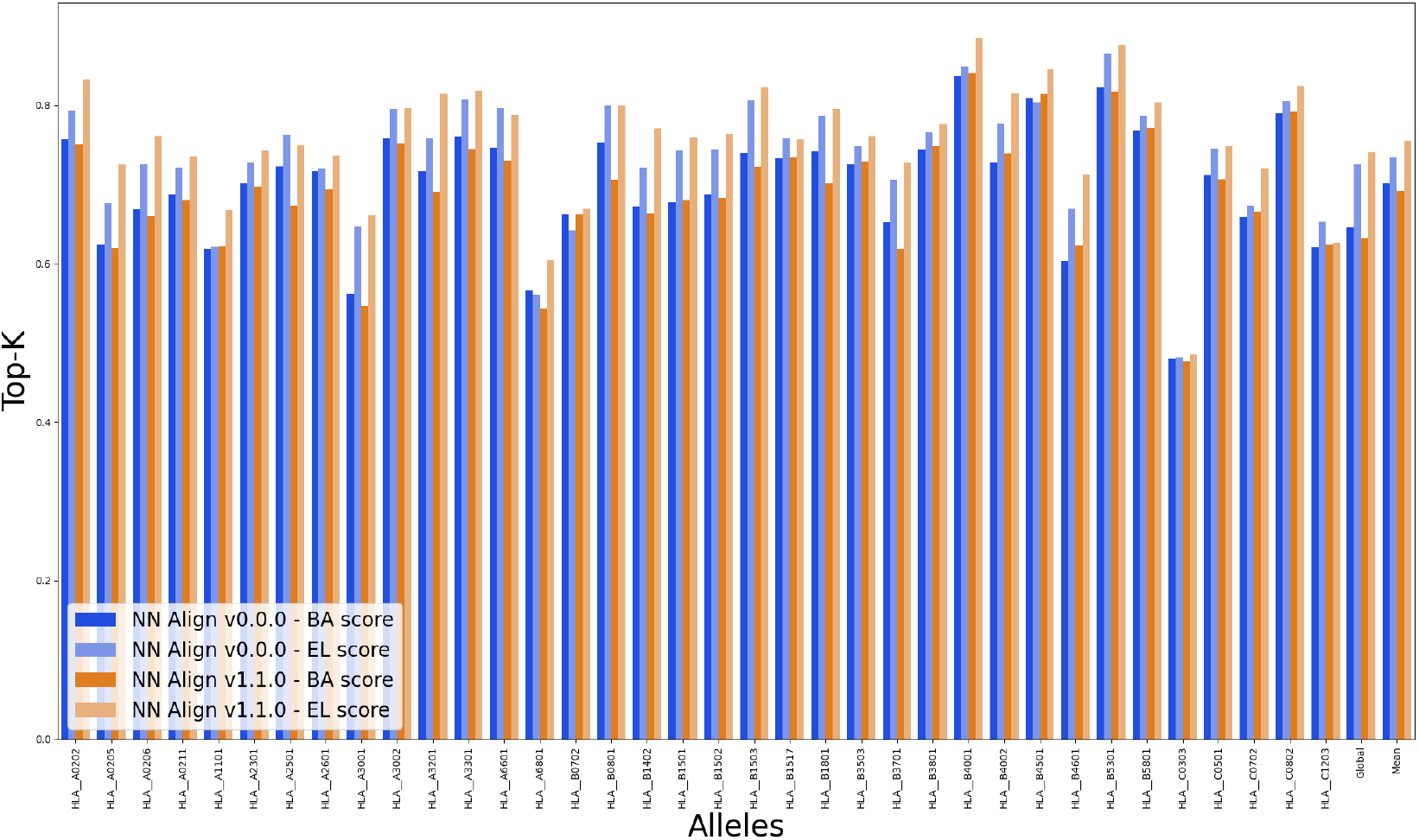
Per-allele Top-K on the regenerated MS ligands evaluation set (`v2.0.0`). For each allele, the four bars show the BA-score and EL-score heads of the `v0.0.0` ensemble (blue / light blue) and of the `v1.1.0` ensemble (orange / light orange). The two ensembles share architecture, optimiser, schedule, ensemble size and random seeds; only the training data differs. The two right-most clusters are the global Top-K pooled over all peptides (`Global`) and the per-allele mean (`Mean`).

## D Inference speed

Table 4 reports end-to-end wall-clock time to score a fixed reference set of 280,000 peptide/allele pairs (lengths 8 to 14, 155 alleles, balanced across both) with the full 50-model **BA** + **SA** + **MA** ensemble, measured identically for both frameworks (feature preparation, all forward passes, post-processing and the CSV write). The fused path’s scores agree with the per-member path to within about 10^−6^; only the reported majority core can differ, and only on rare `argmax` ties. Figure 11 plots per-peptide time on a log scale.

**Table 4:** End-to-end wall-clock to score 280,000 peptide/allele pairs (8 to 14mers, 155 alleles) with the 50-model BA + SA + MA ensemble, on the same P100 VM. The *vs ref* column is relative to the NetMHCpan-4.1 binary driven by a 10-process `compute + gather` wrapper. Warm runs; one-time feature preprocessing is cached and amortised across the 50 members.

| Setup | Device | Wall (s) | Peptides/s | vs ref |
| --- | --- | --- | --- | --- |
| NetMHCpan-4.1 binary, 10-proc wrapper | Xeon CPU (16 cores) | 1080 | 259 | 1 × (ref) |
| NetMHCpan-4.1 binary, 1 proc (original) | Xeon CPU (16 cores) | 9543 | 29 | 0.11 × |
| Reproduction (fused) | NVIDIA P100 (CUDA) | **27** | **10 513** | **40.6 ×** |
| Reproduction (fused) | Xeon CPU (16 cores) | 67 | 4186 | 16.2 × |

**Figure 11:**
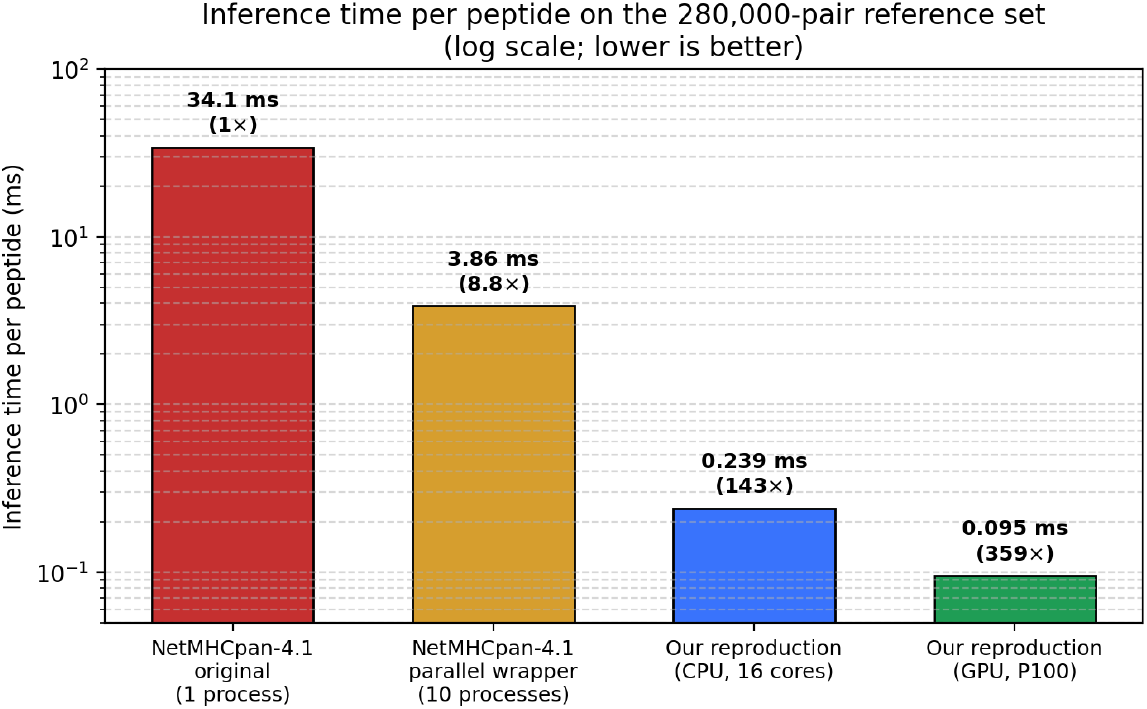
Inference time per peptide on the 280,000-pair reference set (log scale, lower is better); speed-up factors are relative to the original single-process NetMHCpan-4.1 binary. The GPU run scores each peptide in 0.095 ms, about 350 *×* faster than the original binary and about 40 faster than the 10-process wrapper.

## E Dataset versioning

Every dataset, and the model trained on it, carries a semantic`vMAJOR.MINOR.FIX` version: MAJOR increments when hit filtering changes, MINOR when decoys are resampled, and FIX for schema or formatting changes (Table 5). A model takes the version of the data it was trained on, so the numbers in Table 2 can be reproduced by pinning to `data/v0.0.0` or `data/v1.1.0`.

**Table 5:** Dataset versions exposed through the public data bucket. Semantic numbering `X.Y.Z` tracks (X) hit-filtering changes, (Y) decoy-resampling changes, (Z) CSV-schema changes. This paper reports the `v0.0.0` reproduction trained on data `v0.0.0` and the `v1.1.0` re-curation trained on data `v1.1.0`, both evaluated on the original MS ligands set (`v0.0.0`) and on the regenerated MS ligands set (`v2.0.0`).

| Version | Hits | Decoys |
| --- | --- | --- |
| v0.0.0 | *verbatim* NetMHCpan-4.1 hits | *verbatim* NetMHCpan-4.1 decoys |
| v1.0.0 | unmatched hits removed | per-protein sampling, 1:99, cysteine-matched |
| v1.1.0 | same as v1.0.0 | per-protein sampling, **1:17**, cysteine-matched |
| v1.2.0 | same as v1.1.0 | per-protein sampling, 1:20, one source protein per hit |
| v1.3.0 | + BoLA *amani*.1/gb1.7 filter | 1:100 |
| v2.0.0 | + imperfect matches removed ($\sim 7.7\%$) | per-protein sampling, 1:20 |
| v2.1.0 | same as v2.0.0 | per-protein sampling, 1:100 |

## F Reproducibility

The repository contains the `bench_mhc` package (Apache 2.0); the model checkpoints (CC BY-NC-4.0); the YAML training configurations; the pinned Python environment (`pyproject.toml` and `uv.lock`); and a Dockerfile with CUDA support. Per-model documentation pages include training, validation and timing curves for every ensemble described here, plus the `generate-performance-plot` configurations needed to regenerate every figure from the published metrics CSVs. A public object-storage bucket mirrors every dataset version (Appendix E) and every checkpoint. The repository is at https://github.com/instadeepai/BenchMHC-public; the bucket and the documentation are linked from it.

## Notes

https://github.com/instadeepai/BenchMHC-public

### Competing Interest Statement

The authors were affiliated with InstaDeep or BioNTech during the conduct of this research. Both organizations operate in the fields of artificial intelligence and biotechnology. The authors declare no other competing interests.

